# Equilibrium dynamics of the pre- and post-cleavage regions of SP1 are separately shifted by the HIV-1 maturation inhibitor Bevirimat

**DOI:** 10.1101/2022.02.05.479246

**Authors:** Chaoyi Xu, Juan R. Perilla

## Abstract

Human immunodeficiency virus type 1 (HIV-1) assembly and maturation proceeds in two distinct steps. During assembly, viral Gag oligomerizes into a hexagonal polyprotein lattice incorporating the capsid protein (CA) and spacer peptide 1 (SP1) domains, that constitute the immature Gag lattice. During maturation, CTD-SP1 hexamers formed in the previous step are cleaved by HIV-1 protease, causing a dramatic rearrangement of the immature virion to its mature, infectious form. The first-generation maturation inhibitor (MI) bevirimat (BVM) is reported to block the final cleavage between CA and SP1, thus blocking HIV maturation. In contrast, the host factor inositol hexakisphosphate (IP_6_) is a co-factor of Gag assembly and facilitates the formation of a quaternary arrangement of SP1 known as the six helix bundle (6HB). Here, starting from a MAS NMR structure and using atomistic free energy calculations, we establish that binding of BVM and IP_6_ to the immature lattice lacks any cooperativity or avidity. Furthermore, we rationalize the molecular origin of HIV resistance to BVM by determining the role of BVM on the stability of the 6HB and by revealing that SP1 shows independent dynamics for its pre- and post-cleavage regions. Finally, results from our simulations permit us to propose a novel chemical scaffold for the design of maturation inhibitors based on BVM and IP_6_.

## Introduction

During the late phase of the HIV life cycle, Gag, a polyprotein that contains subdomains corresponding to the matrix (MA), capsid (CA), and nucleocapsid (NC) proteins, as well as two spacer peptides (SP1 and SP2) (Fig. 1a), oligomerizes into an immature lattice at the plasma membrane; the latter drives assembly and budding of the virus particle (*1* –*3*). Once budded from the cell surface, the immature virion (Fig. 1b) undergoes maturation, a process that is triggered by proteolytic cleavage of Gag by HIV protease (Fig. 1a). Maturation is marked by a structural rearrangement of the internal constituents of the virion, formation of a mature capsid composed of CA, and activation of viral infectivity (*4* –*6*). Consequently, Gag proteolytic cleavage is an essential step conducive to HIV infectivity and is a prime target for therapeutic development.

**Figure 1:**
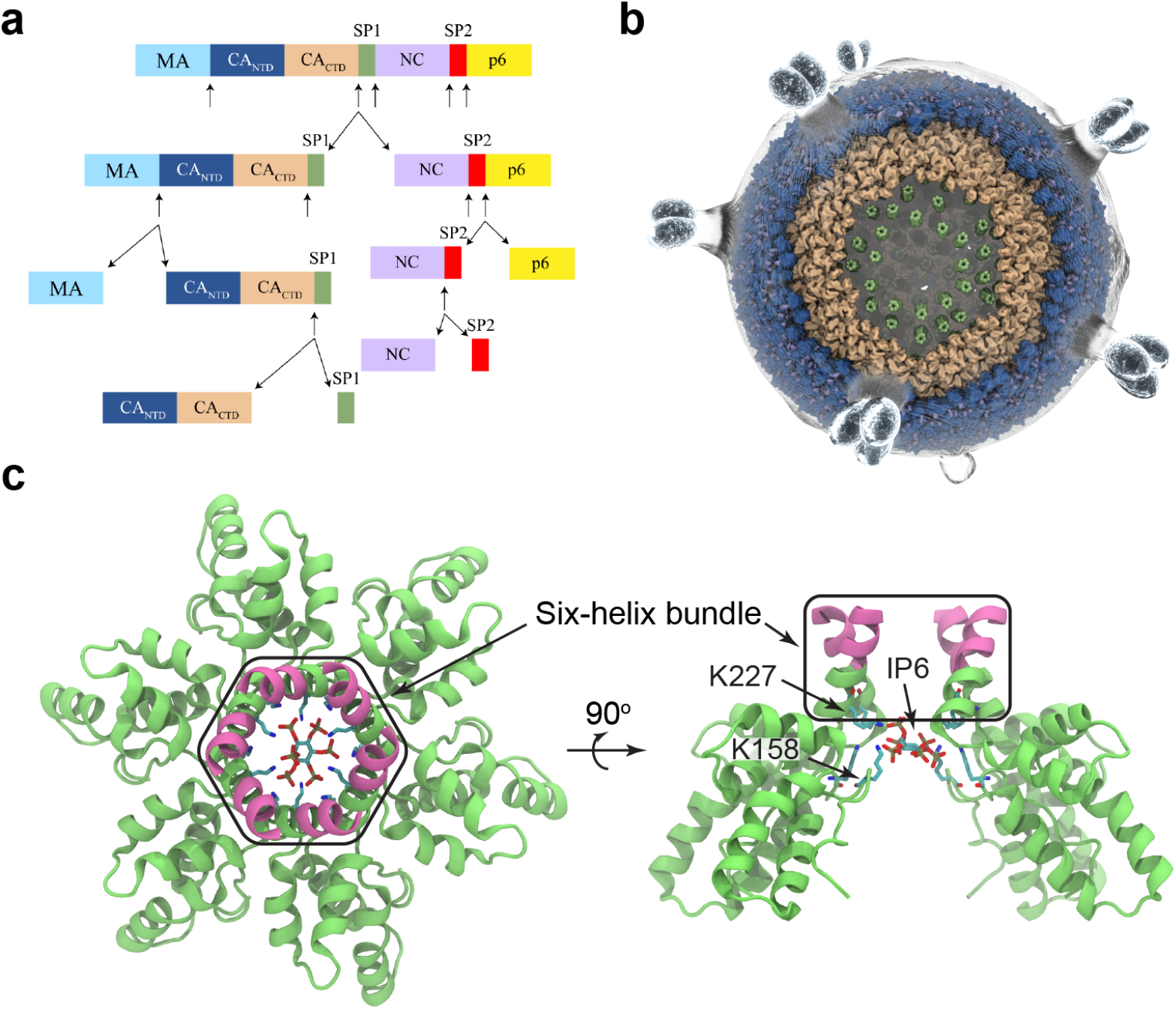
HIV-1 Gag in the late stage of HIV-1 life cycle. (a) A schematic of sequential viral cleavages of the HIV-1 Gag protein. The color blocks represent different domains in the Gag protein and the cleavage sites are indicated by arrows. The light blue, dark blue, golden, green, purple, red and yellow colors represents the MA, CA_NTD_, CA_CTD_, SP1, NC, SP2 and p6 domains in HIV-1 Gag. (b) Atomistic model of an immature HIV-virion, highlighting the different domains of Gag. (c) Solid-state NMR derived structure of a planar CTD-SP1 construct in complex with IP_6_ (PDBID: 7R7P).

The CA-SP1 junction has been shown to act as both an assembly switch and a maturation switch (*7* –*9*) and that the SP1 peptide and the seven CA domain residues that precede it can fold into a helix that form together a six-helix bundle (6HB) in a Gag immature hexamer (illustrated in Fig. 1c). Available structures of HIV-1 CTD-SP1 hexamers and Gag (*10* –*12*) reveal two lysine rings located in the center of the hexamer. Negatively charged metabolites, specifically inositol phosphates (IPs), have been reported to be HIV-1 capsid assembly cofactors by alleviating lysine-lysine electrostatic repulsion at the center of Gag hexamers (*13, 14*).

Cleavage of CA-SP1 is the last step of Gag sequential proteolytic processing (*15, 16*). Maturation inhibitors (MIs) have been developed to target the CA-SP1 cleavage site and inhibit HIV-1 maturation (*17*). So called 3-O-(3,,3,-dimethylsuccinyl) betulinic acid (*18, 19*), also known as bevirimat or simply BVM, is the first generation HIV-1 MI with an *in vitro* IC_50_ of 10 nM (*20*). Evidence points to BVM possessing a novel mechanism of action: it could stabilize the immature Gag lattice (*21, 22*) and prevents cleavage of SP1 from the C-terminus of CA (*9, 18, 20, 23*), which results in non infective HIV virions.

HIV resistance to BVM is gained by the introduction of single-point mutations in the so-called six helix bundle region(6HB) (*24* –*26*), thus indicating a plausible site for binding of BVM. The molecular interactions of BVM with the 6HB have been identified by micro electron diffraction (ED) (*27*). We recently derived from solid-state NMR (ssNMR) distance restraints a clearer picture of the binding of BVM, revealing that it is located in close proximity to the binding site of IP_6_ (PDBIDs: 7R7P and 7R7Q). Here, starting from our ssNMR-derived structure, we have performed all-atom MD simulations (*28, 28* –*30*) and free energy calculations (*31*) of CTD-SP1 in complex with both BVM and IP_6_; in total our simulations sampled over 100 *µ*s of chemical time (Supplementary Table 1). Similar to previous studiess (*9, 23*) we calculated CGENFF-derived parameters for BVM and encountered during validation of the parameters marked difference to the quantum mechanical potential energy surfaces. To address the later issue, we derived CHARMM optimized parameters for all charges, bonds, angles and dihedrals angles parameters in BVM, such that these parameters accurately reproduced the QM potential energy surface. In addition, to reproducing previous results showing stability of the symmetrical nature of the 6HB (*23*), results from our simulations reveal the inhibitory mechanism of BVM in the presence of cellular metabolites in the context of SP1 helix-coil transition path ensembles. Furthermore, by probing the structure-activity relationships of BVM as a maturation inhibitor, we are able to propose a novel chemical scaffold for the molecular design of a novel class of HIV-1 maturation inhibitors and provide a proof-of-concept molecule by evaluating the stability of the complex by utilizing PIP3 lipds.

## Results and discussion

All-atom molecular dynamics (MD) simulations are routinely used to probe protein-ligand and protein-protein interactions (*28* –*30, 32* –*34*). Molecular mechanics (MM) force fields (FF) are essential to MD simulations, and the reliability of the simulations depends heavily on the accuracy of the MM parameters adopted (*35*). To the best of our knowledge, accurate MM parameters for BVM were unavailable at the time of this study. Therefore, we derived BVM CHARMM-compatible parameters by analogy, utilizing the CHARMM general force-field tool (CGENFF) (*36, 37*). The initial parameters produced by CGENFF resulted in several bonded parameters and charges with a high penalty score (greater than 10; Fig. 2a), indicating that these parameters are inaccurate and require at minimum validation and most likely further optimization. Evaluation of the match between the QM PES and the CGENFF-derived MM surfaces resulted in a large root means squared error, indicating that optimization was required. To optimize the aforementioned parameters, we opted for a fragmentation approach (*38*), where BVM was decomposed into two fragments based on the location of the bad parameters in the molecule, as shown in Fig. 2b. The incorrect parameters for atomic partial charges, bonds, angles, and torsions present in the two fragments were then refined to match the quantum mechanically (QM) derived potentials using FFTK (*39*) and Gaussian09 (*40*). Bad partial charges were tuned to optimize the interaction between the atom of interest and a nearby water molecule (*39*). The bond and angle parameters were optimized for twenty iterations until convergence was achieved (Fig. 2c). Subsequently, the dihedral angle parameters in each fragment yielded agreeable fits against the QM potentials, especially in potential energy regions lower than 10 kcal/mol as shown in Fig. 2d-e. Importantly, these conformational scans highlight the accuracy of the entire set of parameters including charges, bonds, angles and dihedrals present in BVM, as both the MM and QM potentials are evaluated utilizing the entire molecule..

**Figure 2:**
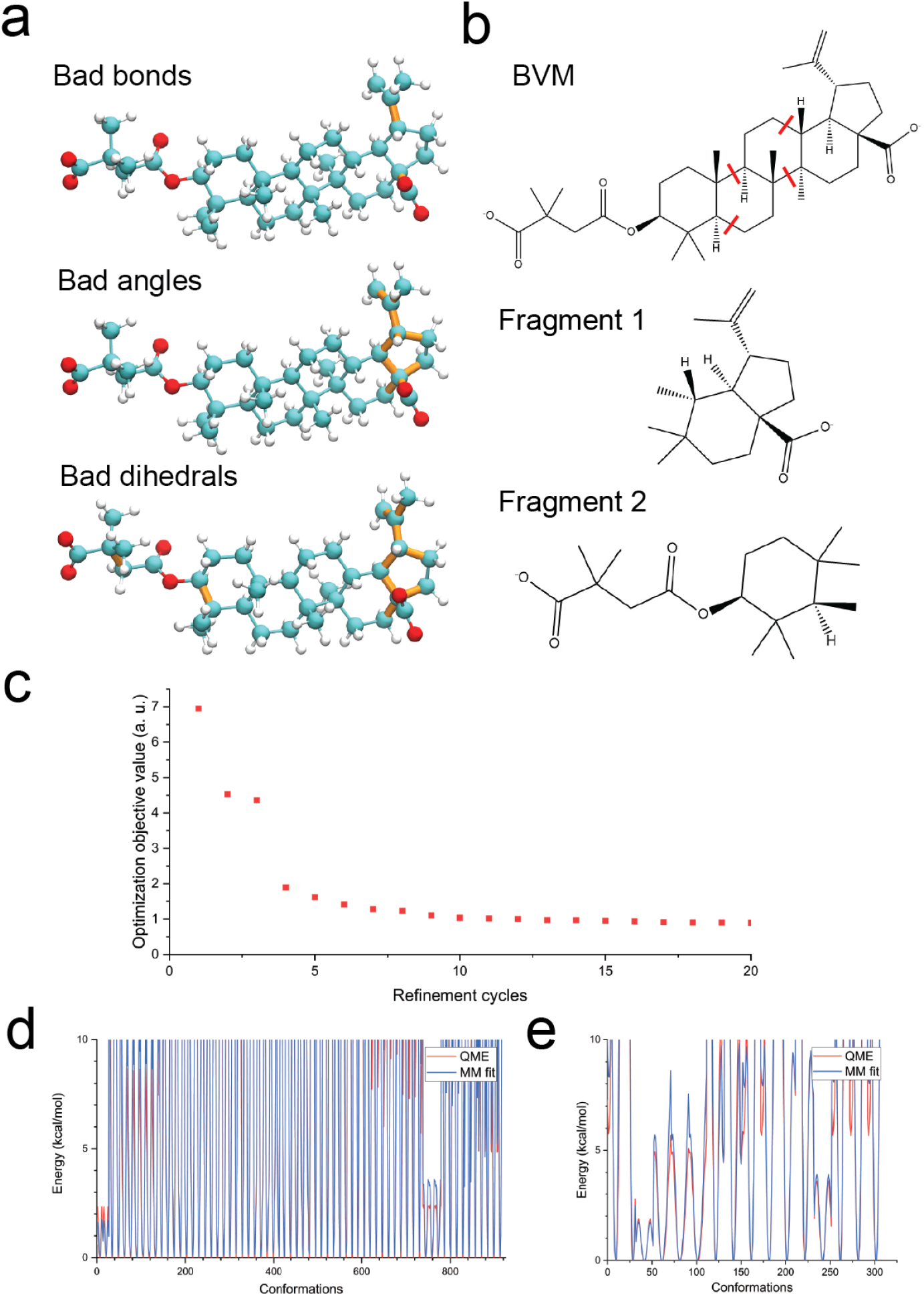
Determination of CHARMM compatible force-field parameters for the HIV-1 maturation inhibitor Bevirimat (BVM). (a) Bond, planar, and dihedral angles with high penalty scores assigned by CGENFF during the initial generation of BVM parameters by analogy. Bonds in orange indicate the bonds, planar and dihedral angles with a parameter penalty score greater than ten. (b) BVM was separated into two fragments taking into consideration the high-penalty scores assigned by CGENFF. Bonds in red represent that were capped with methyl groups at the cut points. (c) Iterative optimizations of the bond and angle parameters for BVM fragment 1 against calculated QM target data. (d)-(e) Classical and quantum potential energy surface profiles of the dihedral angle scans and fitting results for BVM fragment 1 and 2, respectively. These dihedrals scan evaluate the entire force field, namely charges, bonds, angles and dihedrals, indicating that the parameters accurately reproduce the QM potential energy surface.

The QM-optimized parameters for BVM were then used to probe BVM dynamics in explicit solvent (Supplementary Figure 1a). Production simulations with lengths of 500 ns and 100 ns were performed using NAMD2 (*41*) and Anton2 (*42*), respectively. Tracking of the root mean square deviations (RMSD) of the heavy atoms in BVM (Supplementary Figure 1b) indicate that during the simulations, their displacements respect to the QM optimized structure are below 1.5 Å, thus confirming the good agreement between the MM and QM potentials. However, by performing structural clustering, we were able to identify two major BVM conformers (*43*), as illustrated in Supplementary Figure 1b. The major structural differences between these two clusters arise from the stereochemistry of the dimethylsuccinyl carboxylate moiety – characterized by one of the torsion parameters optimized in our study. In contrast, the pentacyclic triterpenoid moiety of these two clusters only exhibits minuscule differences. This finding is also supported by root mean square fluctuation (RMSF) calculations (Fig. 2c), as the dimethylsuccinyl carboxylate moiety in BVM exhibits higher flexibility in two independent simulations. Long-range interactions between BVM with water and ions are also identically sampled by both NAMD2 and Anton2 (Supplementary Figure 1d-f). These results indicate the consistency of our QM-optimized BVM parameters in NAMD and Anton2 simulations. Overall, the simulation results of BVM in a water box further validate and highlight the need for high quality parameters to perform atomistic molecular dynamics simulations that include BVM.

As mentioned before, micro-ED structures show that BVM binds within the 6HB of the CTD-SP1 hexamer, however limits in resolution obscure details regarding molecular interactions between BVM and SP1 (*9, 27*); for instance, the relative orientation of BVM with respect to the 6HB is not fully resolved in the electron maps. Previously, utilizing CGENFF-derived parameters we (*9*) and others (*23*) have derived models of CTD-SP1 and BVM complexes. Here, we extended our previous calculations by using free energy calculations employing the Hamiltonian replica exchange umbrella sampling (HREX/US) to investigate the energetics of BVM binding to the 6HB, and thus determining molecular details of their interactions. Our approach is significantly different from previous efforts (*9, 23*) as we sample all possible binding motifs of BVM to the pore at the center of the 6HB. The binding site in the 6HB can be conceptualized as a cylinder aligned along a CTD-SP1 hexamer (Fig. 1c). Since the pentacyclic triterpenoid moiety of BVM is quite rigid (Fig. 2c) one can envision two orientations of BVM inside of the cylindrical 6HB (Fig. 3a). However, the most likely orientation of BVM with respect to the 6HB was established by computing the relative binding free energies for both orientations (Fig. 3b). Specifically, relative to the solvation energy of BVM alone, binding of BVM in orientation 1 (−23 kcal/mol) is more energetically favorable than in orientation 2 (−5 kcal/mol); these results are in good agreement with previous studies (*9, 23*) the solid-state NMR distance-restraints and therefore unambiguously determine the binding mode of BVM to CTD-SP1 hexamers.

**Figure 3:**
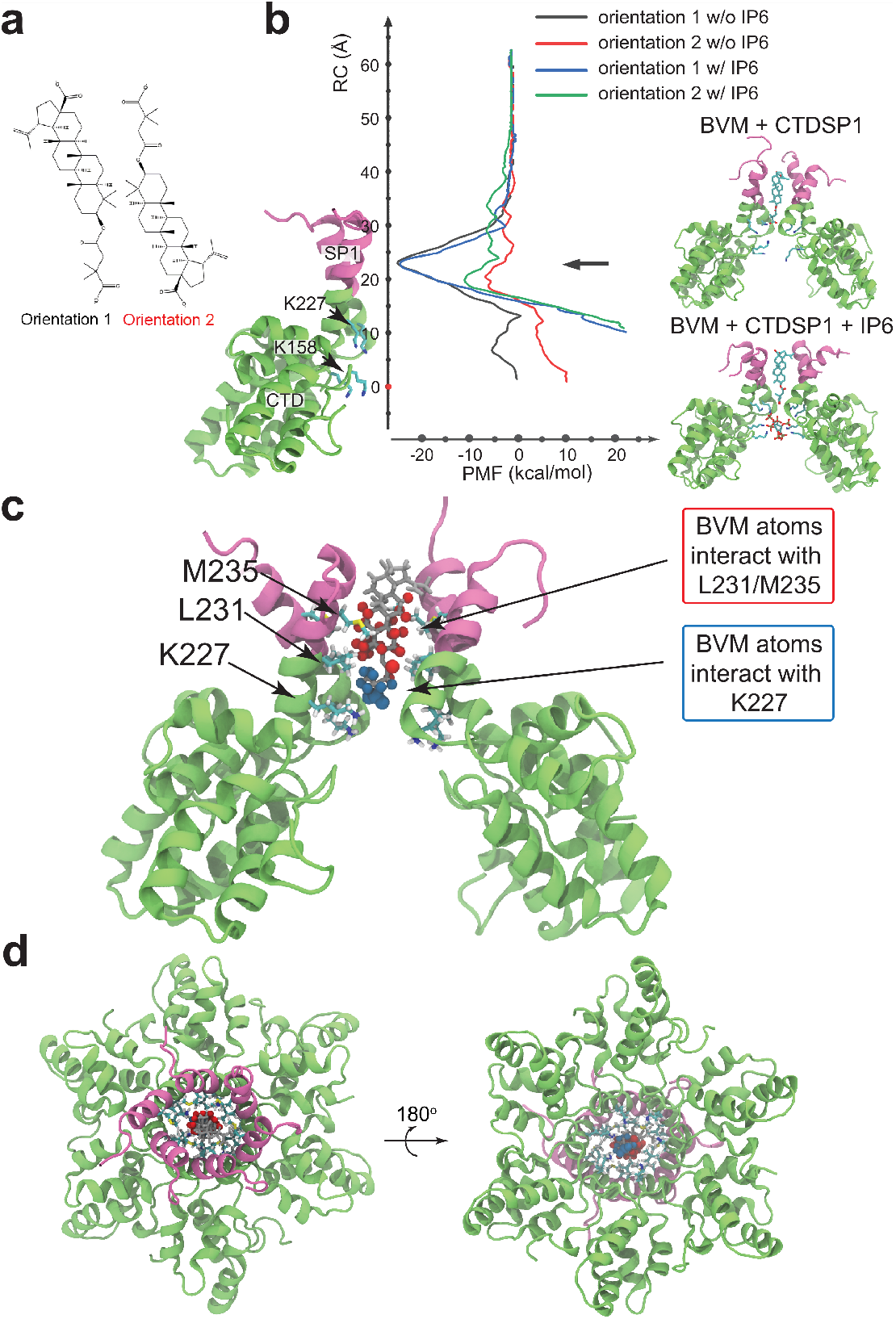
Molecular interactions between BVM and the HIV-1 CTD-SP1 hexamer from free energy calculations. (a) Definition of the two BVM orientations with respect to the CTD-SP1 hexamer. In orientation 1 the dimethylsuccinyl carboxylate group is pointing toward CTD-SP1 N-terminus, in orientation 2 the same chemical group is pointing towards CTD-SP1 C-terminus. (b) Potentials of mean force (PMFs) of the interactions between BVM and the CTD-SP1 hexamer, for two different orientations, with and without IP_6_. Representative structures of the CTD-SP1/BVM and CTD-SP1/BVM/IP_6_ complexes are illustrated in the insets. (c) Detail of the interactions between BVM and the CTD-SP1 hexamer from free energy simulations. Hydrophobic contacts between BVM and L231/M235 are shown in red, while electrostatic interactions between K227 and the dimethylsuccinyl carboxylate moiety of BVM are shown in blue. Occupancies for these interactions are tabulated in Supplementary Table 2 and 3. (d) Top (left) and bottom view (right) of BVM in complex with the CTD-SP1 hexamer shown in (c).

Cooperativity and avidity between IP_6_ and BVM was determined from BVM binding free energies profiles in the presence and absence of IP_6_. Initially, IP_6_ was placed in its canonical binding site (*13*) and positionally restrained to prevent its diffusion during MD simulations. Then, the relative binding free energies of BVM in the presence of IP_6_ were computed as illustrated in Fig. 3b; we observe that BVM binds to 6HB more favorably in orientation 1 in presence of IP_6_. Moreover, the value of the BVM binding energy remains unchanged by the presence of IP_6_ (*<* 1 kcal/mol), as well as the location of BVM’s binding site. However, the PMF significantly increases when BVM approaches IP_6_, indicating a repulsive interaction between the two molecules. In contrast to previous studies (*23*), our free energy calculations demonstrate that IP_6_ has no significant effect on BVM canonical binding within the 6HB and that these two molecules can coexist in the same CTD-SP1 hexamer.

Details regarding the molecular interactions between BVM in complex with a CTD-SP1 hexamer at the identified PMF minimum were probed via contact analysis. Contact occupancies serve as a proxy for the probabilities that any residue of SP1 remains in contact with specific atoms of BVM during the course of the simulation. The most prevalent interactions between BVM and SP1 residues are summarized in Supplementary Tables 2 and 3, and Supplementary Figure 2. From the contacts we determined that major contributors for BVM binding are hydrophobic interactions with L213 and M235 (Fig. 3c,d). Electrostatic interactions between BVM and SP1 mostly come from interactions between K227 and the dimethylsuccinyl carboxylate moiety (Fig. 3c,d), however these electrostatic contacts are not as stable as the hydrophobic interaction according to the differences in the values between their contact occupancies. Furthermore, the influence exhorted by IP_6_ on BVM can also be evaluated from our contact analyses. In particular, we found that the majority of the BVM-SP1 contacts with and without IP_6_ are primarily contributed by L231 and M235. The latter, further supports the premise that IP_6_ has no considerable effect on the binding of BVM.

Stability of the 6HB was evaluated via canonical MD simulations for 10 *µ*s of the CTD-SP1 hexamer in complex with IP_6_ and BVM (Fig. 4). During the simulations, we observed that BVM remains bound within the 6HB while preserving its structural integrity; in addition, similar to previous studies (*23*), the 6HB pseudo six-fold symmetry is maintained during the course of the simulation (Fig. 4a). A collective variable, so-called symmetry index was defined to characterize the pseudo six-fold symmetry of the 6HB during the simulation; the symmetry index is inversely related to the amount of symmetry present in the system (Fig. 4). The distributions of symmetry indices from three different simulation setups are shown in Fig. 4b. Within 2 *µ*s, in the absence of BVM and IP_6_, the 6HB becomes disordered and lacks symmetry (as indicated by its large symmetry index), in contrast the BVM and IP_6_ bound CTD-SP1 systems exhibit the lowest symmetry index, demonstrating the stabilizing effect of BVM on the 6HB. Our results further support the proposed mechanism whereby BVM inhibits maturation by stabilizing the 6HB (*9, 11, 13, 44*). Stabilizations of the 6HB is driven primarily by hydrophobic interactions between BVM with residues L231 and M235 from each monomer. In addition, the dimethylsuccinyl carboxylate group in BVM interacts with residue K227 and anchors BVM within the 6HB.

**Figure 4:**
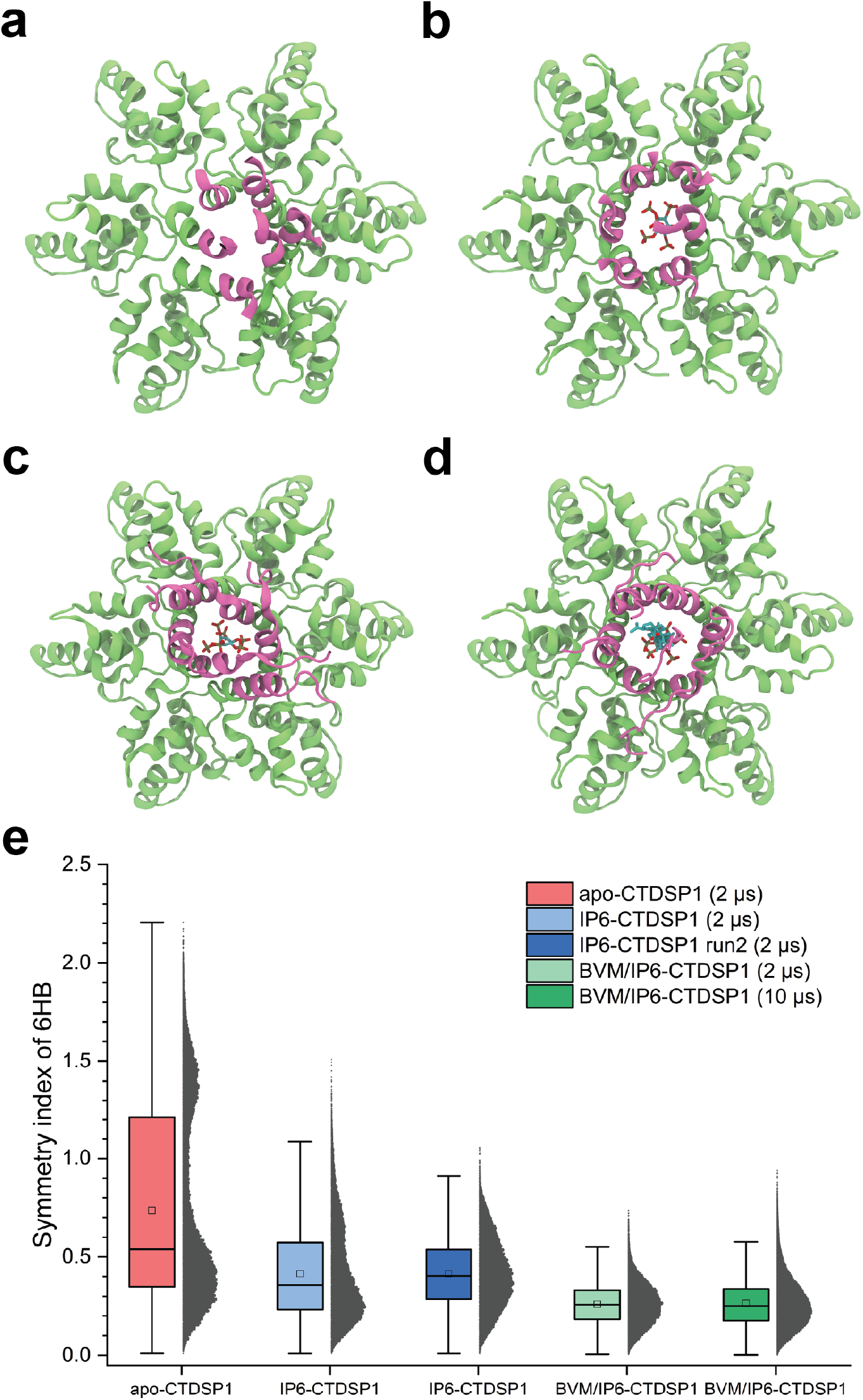
BVM binding stabilizes the six helix bundle present in CTD-SP1 hexamers. (a) Final conformation of a CTD-SP1 hexamer after 2 *µ*s MD simulation. Major quaternary changes of the 6HB are observed after simulation. (b,c) Final conformation after two independent 2 *µ*s MD simulations of CTD-SP1 in complex with IP_6_. In contrast to the apo case, minor changes of the quaternary structure of the 6HB are observed. (d) Final conformation after 10 *µ*s of MD simulation for BVM/IP6 in complex with a CTD-SP1 hexamer. There are no discernible changes of the quaternary structure of the 6HB. (e) Distributions of the symmetry index in apo-, IP_6_-bound and BVM/IP_6_-bound CTDSP1 hexamer. The minimum, first quartile, median, third quartile, and maximum of the distributions are represented as box plots, while the mean of the distributions are highlighted as a square. These distributions show that binding of BVM and IP_6_ have the major stabilization effect on the 6HB.

**Figure 5:**
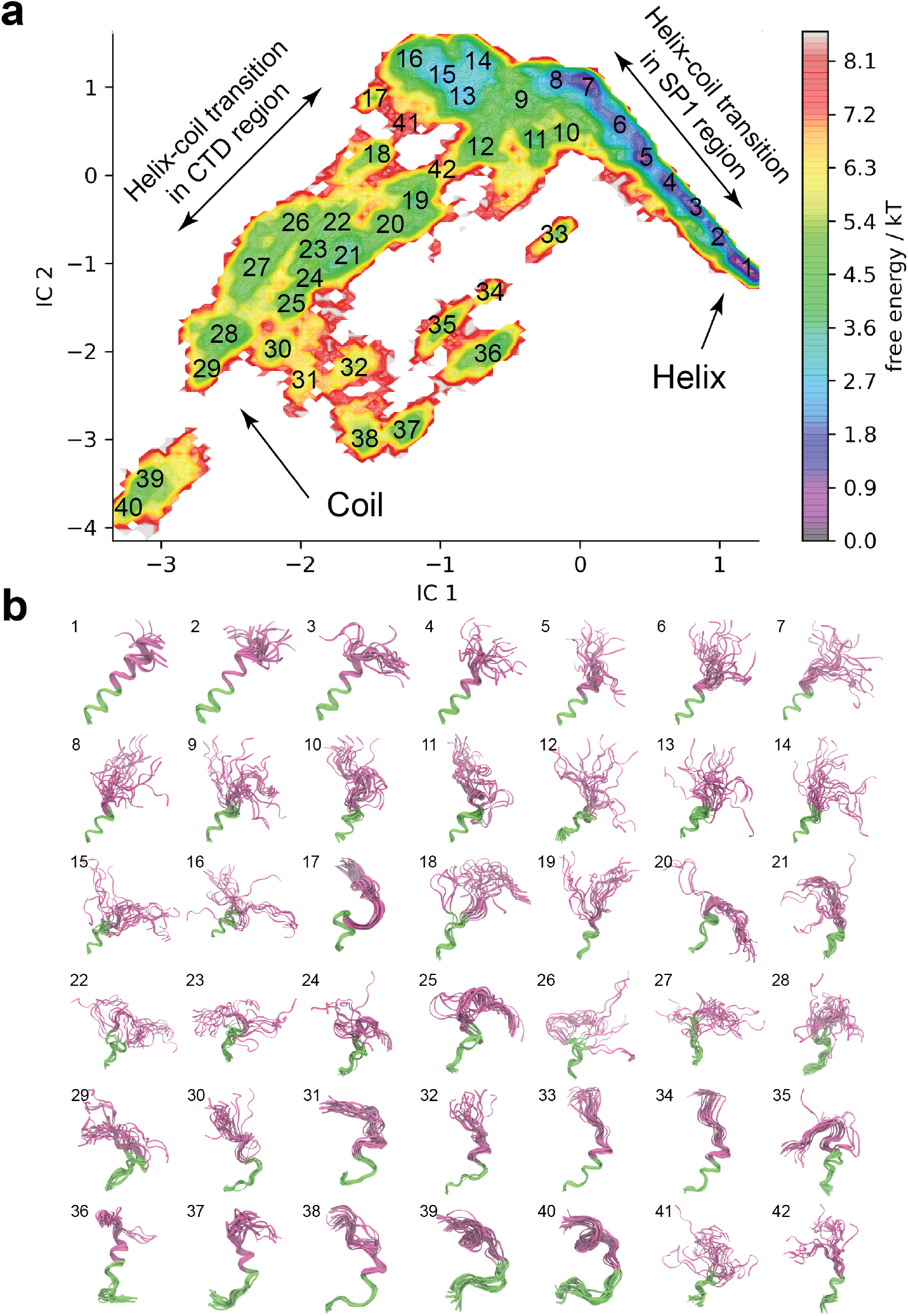
Transition path ensemble (TPE) of SP1 domain in CTDSP1 hexamer. (a) The free energy landscape of SP1 domain helix-coil transition from independent component analysis. (b) TPE derived from the free energy landscape. The numbers on the upper left corner in panel (b) indicate the cluster centers identified in panel (a). The relaxed TPE shows that the helix-coil transition occurs in two stages: folding of the SP1 region, followed by folding of the CTD region. The helix to coil transition path initiates by local unfolding of the SP1 region followed by the unfolding of the CTD region. The coil to helix transition is initiated by the folding of the CTD region of SP1.

Gag assembly likely proceeds from smaller quaternary arrangements (e.g., dimers) towards higher order structures like hexamers, and hexamers of hexamers. In addition, the periphery of the extended lattice is known to include grain boundary defects with varying coordination numbers (*12*). Stabilization by BVM of the quaternary structure of intermediate states of Gag and grain boundary defects were evaluated utilizing canonical MD simulations. Pentameric, tetrameric and trimeric forms of CTD-SP1 oligomers were generated by removing one, two, or three consecutive monomers from the hexamer, respectively (Supplementary Figures 4, 5, 6, 7, and 8). In all simulations, BVM was positioned in the same location as it has been found in CTD-SP1 hexamers. After simulation, BVM was still found bound to the SP1 domain in all cases, however the quaternary structure of trimeric to pentameric forms of CTD-SP1 was found collapsed. In contrast, stabilization of the secondary structure of the SP1 domain was enhanced in all cases by the presence of BVM.

The SP1 domain of Gag is known to be highly dynamic and undergo helix-coil transitions at equilibrium (*9*). Characterizing the dynamics of the SP1 domain helix-coil transition is pivotal to establishing the mechanism of inhibition by BVM. To identify SP1 transition path ensembles (TPE) and thus identify structural intermediates(*32*), the conformational kinetic map of the SP1 domain in assembled CTD-SP1 hexamers was derived(Supplementary Figure 5). For this purpose, by applying a bias along the *α*-helix score (HS) (*45*) in steered MD simulations (SMD) of apo-, IP6-, BVM-and IP6/BVM-CTDSP1 hexamers, we generated SMD-based TPEs of the SP1 domain helix-coil transition. The SMD-based TPEs were then relaxed by subjecting intermediate conformations to unbiased molecular dynamics simulations (Supplementary Figure 5b). The relaxed TPE shows that the helix-coil transitions occurs independently for the CTD and SP1 regions. Notably, the helix to coil transition path initiates by local unfolding of the SP1 region followed by the unfolding of the CTD region. In contrast, the coil to helix transition is initiated by the folding of the CTD region of SP1. Comparison between the TPEs for bound and unbound BVM shows that binding of BVM shifts the TPE for the helix-coil transition on the CTD region while maintaining the TPE in the SP1 region (Supplementary Figure 3). Similarly, binding of IP_6_ alters the TPE of the CTD region while preserving the dynamics of the SP1 region. Altogether, our results indicate that in the 6HB, the helix-coil TPE of the CTD region of the SP1 domain is modulated by IP_6_ and BVM binding while the dynamics of the SP1 subdomain is unaltered by any of these two molecules.

Overall, we have been able to characterize the dynamic behaviour of the 6HB in the presence and absence of IP_6_ and BVM. In particular, we have identified that the CTD and SP1 regions of the 6HB show peculiar dynamic behaviour due to their hydrophilic and hydrophobic characteristics, respectively. We hypothesized that molecules which combine the electrostatic characteristics of IP_6_ and the hydrophobic behaviour of BVM would induce a dynamical shift of the SP1 subdomain (Fig. 6a-b). To test this hypothesis, we introduced phosphatidylinositol (3,4,5)-trisphosphate (PIP3) into our simulations. PIP3 includes an inositol head group which contains three negatively charged phosphate groups and two polyunsaturated tail groups (Fig. 6c-d). In an extended conformation, PIP3 hydrophilic head group and its hydrophobic tail fits well within the CTD-SP1 central channel (Fig. 6d). Structural integrity of the CTD-SP1 hexamer probed during 2 *µ*s long MD simulation shows that, for this time scale, PIP3 is capable of stabilizing the 6HB (Fig. 6e, f). Therefore, we propose that a molecular scaffold that combines BVM and IP_6_ into a single molecule will yield a new class of maturation inhibitors.

**Figure 6:**
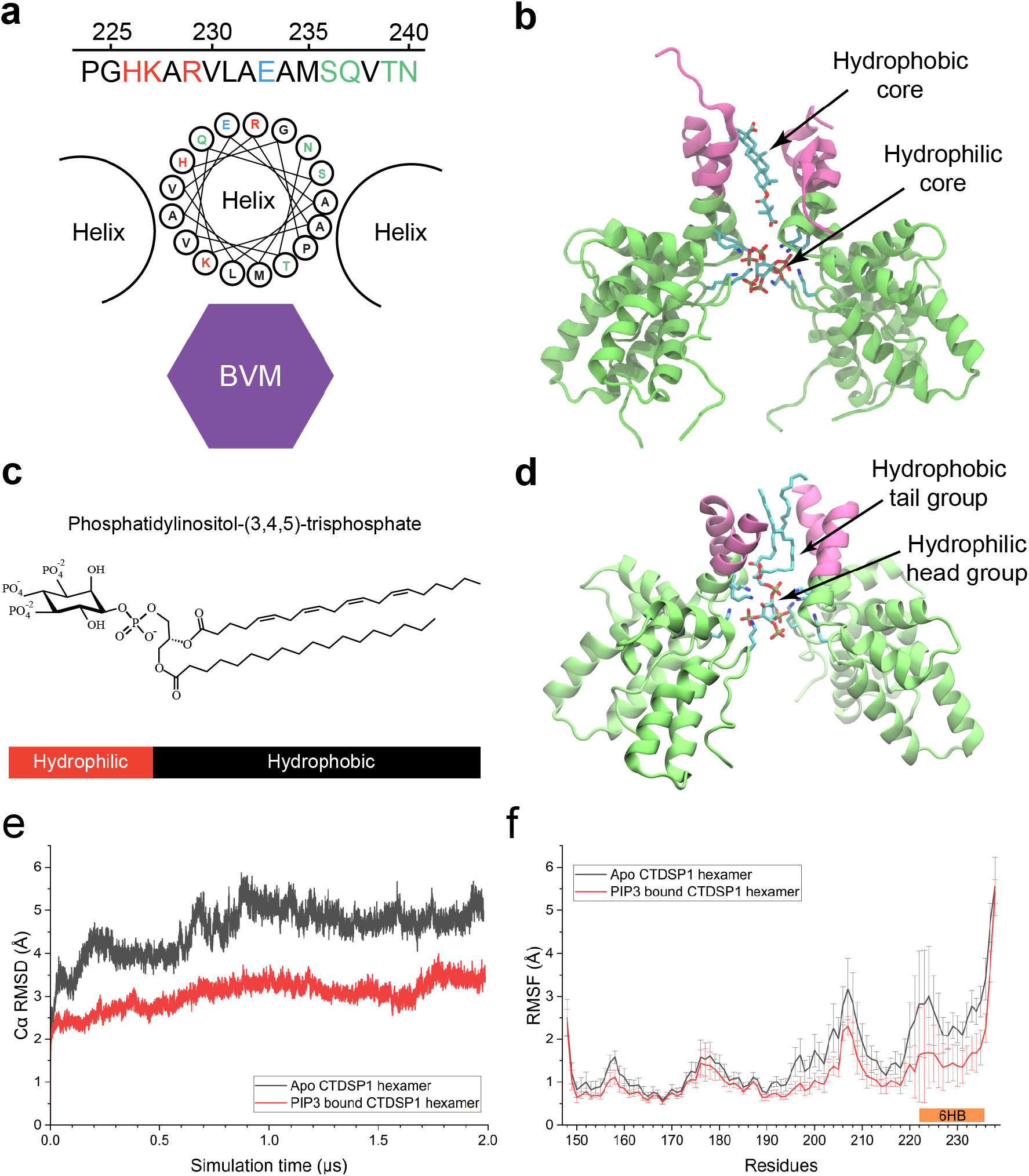
Molecular mechanism of BVM stabilization of HIV-1 CTD-SP1 hexamers. (a) The amino acid sequence and helix wheel representation of the SP1 domain. The amino acids with positively charged sidechains are colored in red, with negatively charged sidechains are colored in blue, with polar and uncharged sidechains are in green, and with hydrophobic sidechains are in black. (b) The BVM and IP6 binding sites in CTD-SP1 are composed of two regions with two opposite hydrophobicities.(c) Molecular structure of phosphatidylinositol-3,4,5-triphosphate (PIP3). (d) Molecular dynamics simulations of PIP3 within the 6HB of HIV-1 CTDSP1 hexamer. (e)-(f) RMSD and RMSF of the Apo- and PIP3-CTDSP1 hexamer simulations.

Maturation inhibitors are unexploited therapeutics and prime chemical probes against HIV. Binding of BVM to the 6HB is characterized by the interactions between six SP1 helices and is coordinated by SP1’s L231 and M235 (Fig. 6a). These hydrophobic residues form a hydrophobic core within the 6HB and form favorable hydrophobic interactions with the pentacyclic triterpenoid moiety of BVM (Fig. 6b). In addition, IP_6_ binds between two lysine rings within the CTDSP1 hexamer. We encountered that BVM shifts the TPE exclusively for the SP1 region, and that IP_6_ is capable of shifting the TPE for the CTD region. That BVM modulates the dynamics of the SP1 region of the 6HB explains why resistance to BVM is characterized by single-point mutations in this region and suggests that such primary sequence substitutions preserve the dynamical behaviour of the SP1 region rather than interfering with binding of BVM. We therefore conclude that design of next-generation maturation inhibitors with combined characteristics of IP_6_ and BVM will reduce the ability of the virus to develop resistance against the portion of these novel molecules that interacts with the SP1 region.

## Methods

### Derivation of molecular mechanics parameters for Bevirimat

CHARMM-compatible molecular mechanics (MM) parameters for BVM (Fig. 2a) were derived by analogy following the CGENFF protocol (*36, 37*). The latter identified several parameters that required validation and manual optimization, therefore partial charges and bonded interactions of BVM (Fig. 2a) that were assigned a penalty greater than ten were refined at quantum mechanics (QM) level using the Force Field Toolkit (*39*) protocol in VMD (*46*). QM calculations were performed utilizing Gaussian09 (*40*) at the MP2/6-31G* level of theory. To generate MM parameters for the entire molecule, we used a previously developed approach based on molecule fragmentation (*38*). First, BVM was divided into two fragments that contain the high penalty regions (Fig. 2b). In each fragment, carbon atoms at the cut points were capped with methyl groups. The molecular structure of each fragment was optimized in Gaussian (*40*). Then, the partial charges of atoms in BVM were optimized to reproduce the hydrogen bond interactions with a water molecule placed nearby. Subsequently, Gaussian calculations of the hessian were used to optimize the parameters of the bonds and angles (*39*) with high penalty scores in the fragment (Fig. 2c). After that, torsion angle scans of the dihedral angles with large penalties were performed to generate QM target data. The dihedral angle parameters were then optimized (Fig. 2d,e) towards the QM target data using the downhill Simplex method (*47*).

Molecular mechanics parameters for BVM were extended for use on the Anton2 supercomputer (*42*). To confirm that results from different MD engines are comparable, we tracked the structural changes and fluctuations of BVM and its interactions with water and ions from independent simulations performed using NAMD2 and Anton2 (Supplementary Figure 1a). As shown in Supplementary Figure 1b-f, the resulting RMSD, RMSF and radial distribution functions from NAMD2 and Anton2 are considerably similar, which indicates the consistency of the BVM simulations on both NAMD2 and Anton2.

### Derivation of molecular mechanics parameters for PIP3

Similar to BVM, the MM parameters for PIP3 were generated by analogy using CGENFF (*36, 37*). The resulting penalty scores were smaller than 10, indicating no need to further optimize using QM calculations, and could be directly used in the MD simulations.

### System preparation and equilibration

An ensemble of five structures from magic angle spinning NMR CTD-SP1 hexamers in complex with IP_6_ and BVM were determined in collaboration with the lab of Tatyana Polenova and are part of a separate manuscript, coordinates are available under PDB accession codes 7R7P and 7R7Q. The ssNMR-derived structure with the lowest energy was minimized and equilibrated using MD while enforcing the NMR derived 32,439 distance restraints and 1,080 torsion restraints utilizing flat-bottom potentials.

After equilibration, BVM and IP6 were placed in the CTDSP1 hexamer to construct CTD-SP1 hexamer models for MD simulations. Four systems were constructed based on the orientations of BVM and the presence of IP6 within the system. BVM was placed in the systems with two opposite orientations. As illustrated in Fig. 3a, orientation 1 of BVM is defined as the direction where the dimethylsuccinyl carboxylate end of BVM is oriented towards the N-terminus of the CTD-SP1 hexamer along the axis perpendicular to the plane of the CTDSP1 lattice; in contrast, in orientation 2, the dimethylsuccinyl carboxylate end of BVM is oriented towards the C-terminus. In the two systems containing IP_6_, IP_6_ was placed in its canonical binding site, specifically between K158 and K227.

Four systems were then subjected to two-stages of minimization, using the conjugated gradient algorithm (*48*) with linear search (*49*) implemented in NAMD2 (*41*). Each minimization stage was performed for 10,000 steps. In the first-stage minimization, only water and ions were free to move, while the protein and IP_6_ were fixed. In the second-stage minimization, heavy atoms in the backbone of the protein were restrained with a force constant of 10 kcal/mol Å^2^. Following minimization, the systems were tempered gradually from 50 to 310 K in increments of 20 K over 1 ns; subsequently, the systems were equilibrated at 310 K for over 20,000 steps. Once the systems were equilibrated, they were used as the initial models for steered molecular dynamics (SMD) simulations (*50*).

MD simulations performed in this study employed NAMD2.14 (*41*) and Anton2 with the CHARMM36m force field (*35*). An integration timestep of 2 fs was employed in the multistep r-RESPA integrator as implemented in NAMD2 (*41*). Bonded interactions were evaluated every 2 fs, and electrostatics were updated every 4 fs. Temperature was held constant at 310 K using the stochastic rescaling thermostat (*51*) with a coupling constant of 0.1 ps^−1^. Pressure was controlled at 1 bar using the Nose-Hoover Langevin piston barostat with a period and decay of 40 ps and 10 ps, respectively. All bonds to hydrogen were constrained with the SHAKE and SETTLE algorithm for the solute and solvent, respectively. Longrange electrostatics was calculated using the particle-mesh-Ewald summation with a grid size of 1 Å and a cutoff for short-range electrostatics interactions of 12 Å.

### Derivation of potential of mean forces

Potentials of mean force between BVM and CTD-SP1 hexamers were derived by employing the so-called Hamiltonian replica exchange/umbrella sampling (HREX/US) method (*31, 52*). A reaction coordinate *ζ* (RC; Fig. 3b) was chosen to defined the position of BVM on the 6-fold rotation axis of the CTD-SP1 hexamer, which can be calculated by 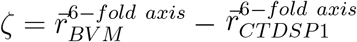, where 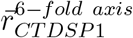 and 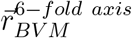 are the coordinates for the center of mass of the CTD-SP1 hexamer and BVM on the RC, respectively. The initial conformations along the RC were generated by performing ensemble-based constant-velocity steered MD (SMD) simulations. During SMD, BVM was steered through the central cavity of the CTD-SP1 hexamer, at a rate of 0.1 nm/ns, along the RC. Eighty snapshots from the SMD simulations were chosen at a constant interval of 0.75 Å on RC and used for HREX/US simulations as the starting conformations at each US window. During the HREX/US simulations, the center of mass of BVM was restrained at each US window with a harmonic potential of 2.5 kcal/mol Å^2^ and exchanges between neighboring US replicas were attempted every 1,000 steps. The exchange acceptance probability *p* between neighboring replicas was obtained using the Metropolis-Hasting criterion (*53*):

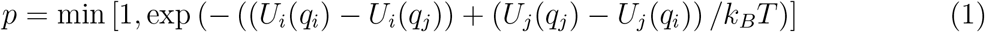

Where T is the temperature of the system, 310 K, *k*_*B*_ is the Boltzmann constant, *q*_*i*_ and *q*_*j*_ represent the conformations of replica in US window *i* and replica in US window*j*, and *U*_*i,j*_ represent the potential part of the Hamiltonian evaluated at the indicated conformation *q*_*i,j*_. After simulation, PMFs were derived using the weighted histogram analysis method (WHAM) (*54, 55*). HREX/US simulations were performed for 30 ns per window and data resulting from the first 10 ns were removed for PMF calculations. The convergence of the PMF results was evaluated and confirmed by the fact that no significant changes in PMF were observed as additional sampling was collected.

### Contact analysis of CASP1-BVM interactions

The contact analysis between CTD-SP1 and BVM was performed in VMD (*46*). Contacts are defined as the distance between the CTD-SP1 residue and BVM with a threshold of 30 Å. Contact occupancies were calculated using the expression

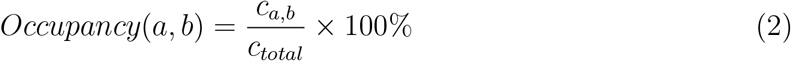

*c*_*a,b*_ is the number of frames in the simulation, where the CTD-SP1 residue *a* and BVM atom *b* form a contact, *c*_*total*_ is the total number of frames of the simulation.

### Anton 2 simulations of BVM/IP_6_-CTD-SP1 hexamer

A system of CTD-SP1 in complex with BVM and IP_6_ was constructed based on our ssNMR-derived CTD-SP1 structure (Fig. 1c). BVM in the system was placed inside the 6HB with the orientation 1 (Fig. 3a). This system was solvated and ionized with the standard protocol. After that, this prepared BVM-IP6-CTDSP1 system was subjected to a two-step minimization, heating and equilibration using NAMD2.14 (*41*), while positional constraints on the backbone atoms were applied. Subsequently, the equilibrated system ran for 10 *µ*s on the PSC Anton2 supercomputer. The CHARMM 36m (*35*) force-field was used for all simulations. During the simulation, the temperature (310 K) and pressure (1 atm) were maintained using the Multigrator integrator (*56*) and the simulation time-step was set to 2.5 fs/step, with short-range forces evaluated at every time step, and long-range electrostatics evaluated at every second time step. Short-range non-bonded interactions were truncated at 17 Å; long range electrostatics were calculated using the *k*-Gaussian split Ewald method (*57*).

### Definition of symmetry index in 6HB

A collective variable, named symmetry index, was defined to evaluate the six-fold symmetry in the 6HB of HIV-1 CTDSP1. The rotation matrix used to describe the rotation operation around the *Z* axis is

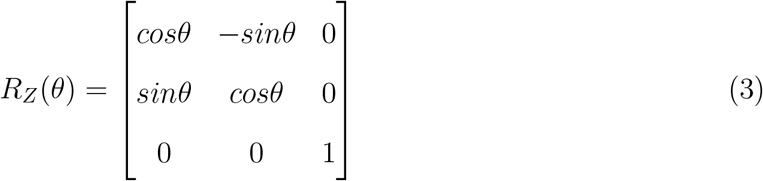

For a 6-fold rotation around the *Z* axe, the rotation matrix is

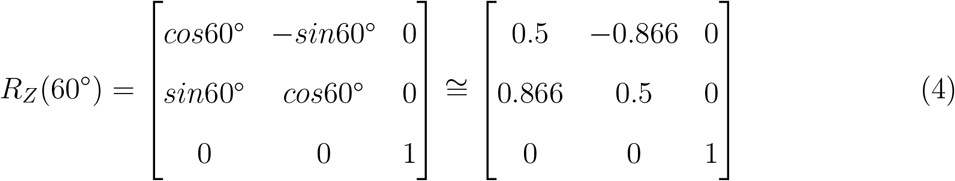

And the rotation matrix between neighboring helices in 6HB is labelled 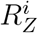, where *i* = 1, 2, 3, …, 6 represents the six possible rotation operations. The symmetry index is defined as the Frobenius norm of the difference between *R*_*Z*_(60^°^) and 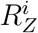, which is expressed as

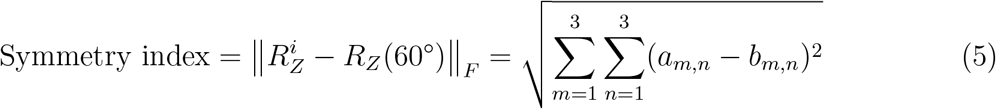

where *m* and *n* are the row number and column number of the symmetry matrices, and *a*_*m,n*_ and *b*_*m,n*_ are the elements in 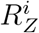, and *R*_*Z*_(60^°^), respectively.

### Determination of transition path ensembles for the SP1 subdomain

Initial transition path ensembles between the helical form of SP1 and its coil form were generated using an ensemble of steered molecular dynamics (SMD) simulations (*50, 58*). The *α*-helix score (HS) in Colvars module (*45*) was used as a descriptor to quantify the progress of the helix-coil transition. Residues 224 to 240 in a randomly selected chain, corresponding to the SP1 domain, among six chains were used for SMD. The initial *α*-helix score of this sequence segment was 0.85 HS. In total, four systems were constructed and equilibrated for SMD simulations, including apo-, IP6-bound, BVM-bound, and IP6/BVM bound HIV-1 CTDSP1 hexamer. The initial conformation of HIV-1 CTDSP1 hexamer was derived from MAS NMR distance restraints. IP_6_ and BVM were placed in their canonical binding sites in the CTDSP1 hexamers. After equilibration, an ensemble of ten independent SMD simulations of each system was performed. In each SMD simulation, residues 224 to 240 were steered to follow a transition from helix to coil for 100 ns with a constant speed of 0.0085 HS per ns. The force constant used in these SMD simulations was 500 kcal/mol HS^2^. Time-lagged independent component analysis (TICA) (*59*) in PyEMMA (*60*) was employed to obtain the conformational dynamics of the SP1 domain from the SMD trajectories. The trajectories from 4*×*10 SMD simulations were collected and pair-wise distances between backbone atoms in the SP1 domain were used as features. Subsequently, TICA analysis was performed to reduce the dimensionality of the SP1 domain conformational dynamics to 100 dimensions. Transition path ensembles from all SMD trajectories were obtained by classifying the trajectory into 100 cluster centers in the space spanned by the independent components. The cluster centers were then used as initial conformations for relaxation of the TPE via unbiased MD simulations. After generation of the initial seeds, 4*×*100 unbiased MD simulations were performed for 20 ns. The first 5 ns of simulation were considered as equilibration and discarded for the following analyses. The last 15 ns data from all unbiased simulations were then used for another round of TICA analysis using the same parameters described above. The relaxed TPE was obtained by classification into structural clusters of the projection of the trajectories onto the two main ICs.

## Acknowledgement

The authors thank Dr. Hamed Meshkin for critical reading of the manuscript. The authors acknowledge funding from the US National Institutes of Health award P50AI150481 and P20GM104316 (to J.R.P.). This work used the Extreme Science and Engineering Discovery Environment, which is supported by the National Science Foundation (Grant ACI-1548562). This work used XSEDE Bridges and Stampede2 at the Pittsburgh Super Computing Center and Texas Advanced Computing Center, respectively, through allocation MCB170096. Anton2 computer time was provided by the Pittsburgh Supercomputing Center (PSC) through Grant R01-GM116961 from the National Institutes of Health. The Anton2 machine at PSC was generously made available by D.E. Shaw Research.

## Supplementary Information

**Table 1:**
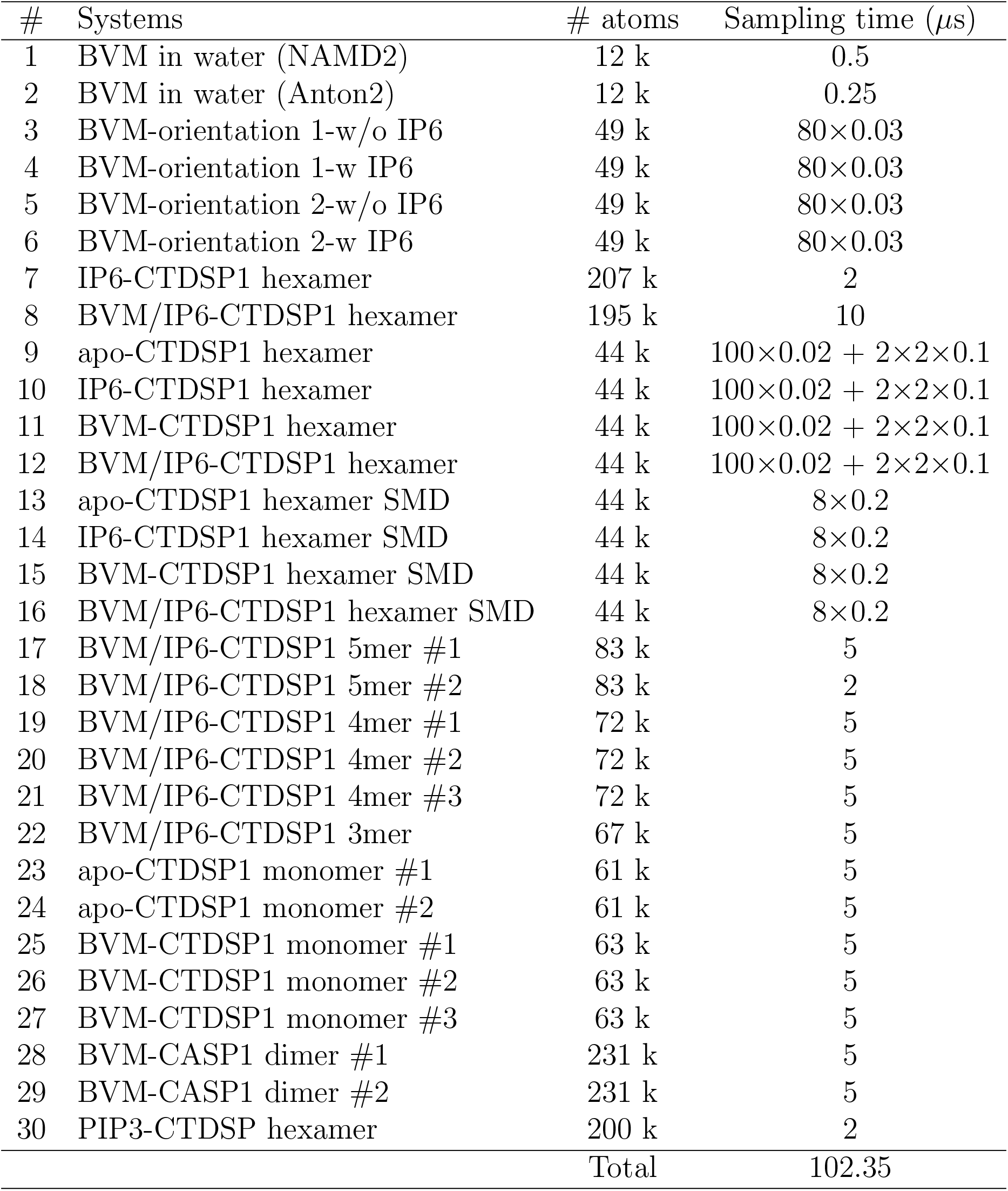
Molecular dynamics simulations performed in this study.

| # | Systems | # atoms | Sampling time (μs) |
| --- | --- | --- | --- |
| 1 | BVM in water (NAMD2) | 12 k | 0.5 |
| 2 | BVM in water (Anton2) | 12 k | 0.25 |
| 3 | BVM-orientation 1-w/o IP6 | 49 k | $80 \times 0.03$ |
| 4 | BVM-orientation 1-w IP6 | 49 k | $80 \times 0.03$ |
| 5 | BVM-orientation 2-w/o IP6 | 49 k | $80 \times 0.03$ |
| 6 | BVM-orientation 2-w IP6 | 49 k | $80 \times 0.03$ |
| 7 | IP6-CTDSP1 hexamer | 207 k | 2 |
| 8 | BVM/IP6-CTDSP1 hexamer | 195 k | 10 |
| 9 | apo-CTDSP1 hexamer | 44 k | $100 \times 0.02 + 2 \times 2 \times 0.1$ |
| 10 | IP6-CTDSP1 hexamer | 44 k | $100 \times 0.02 + 2 \times 2 \times 0.1$ |
| 11 | BVM-CTDSP1 hexamer | 44 k | $100 \times 0.02 + 2 \times 2 \times 0.1$ |
| 12 | BVM/IP6-CTDSP1 hexamer | 44 k | $100 \times 0.02 + 2 \times 2 \times 0.1$ |
| 13 | apo-CTDSP1 hexamer SMD | 44 k | $8 \times 0.2$ |
| 14 | IP6-CTDSP1 hexamer SMD | 44 k | $8 \times 0.2$ |
| 15 | BVM-CTDSP1 hexamer SMD | 44 k | $8 \times 0.2$ |
| 16 | BVM/IP6-CTDSP1 hexamer SMD | 44 k | $8 \times 0.2$ |
| 17 | BVM/IP6-CTDSP1 5mer #1 | 83 k | 5 |
| 18 | BVM/IP6-CTDSP1 5mer #2 | 83 k | 2 |
| 19 | BVM/IP6-CTDSP1 4mer #1 | 72 k | 5 |
| 20 | BVM/IP6-CTDSP1 4mer #2 | 72 k | 5 |
| 21 | BVM/IP6-CTDSP1 4mer #3 | 72 k | 5 |
| 22 | BVM/IP6-CTDSP1 3mer | 67 k | 5 |
| 23 | apo-CTDSP1 monomer #1 | 61 k | 5 |
| 24 | apo-CTDSP1 monomer #2 | 61 k | 5 |
| 25 | BVM-CTDSP1 monomer #1 | 63 k | 5 |
| 26 | BVM-CTDSP1 monomer #2 | 63 k | 5 |
| 27 | BVM-CTDSP1 monomer #3 | 63 k | 5 |
| 28 | BVM-CASP1 dimer #1 | 231 k | 5 |
| 29 | BVM-CASP1 dimer #2 | 231 k | 5 |
| 30 | PIP3-CTDSP hexamer | 200 k | 2 |
|  |  | Total | 102.35 |

**Table 2:** Top 20 BVM/CTD-SP1 molecular interactions with and without IP_6_. The contact occupancy measures how frequently the corresponding contact was formed in the simulation. A contact is defined when the distance between two groups of atoms is smaller than 3.0 Å.

| BVM-CTDSP1 interactions w/o IP<sub>6</sub> |  |  |  | BVM-CTDSP1 interactions w/ IP<sub>6</sub> |  |  |
| --- | --- | --- | --- | --- | --- | --- |
|  | CA residues | BVM atoms | Occupancies (%) | CA residues | BVM atoms | Occupancies (%) |
| 1 | LEU231 | H35 | 97.8 | LEU231 | H35 | 99.6 |
| 2 | MET235 | H7 | 96.2 | LEU231 | H34 | 96.3 |
| 3 | MET235 | H6 | 95.6 | MET235 | H7 | 92.4 |
| 4 | LEU231 | H34 | 89.8 | LEU231 | O4 | 88.7 |
| 5 | LEU231 | O4 | 87.8 | LEU231 | H40 | 88.5 |
| 6 | LEU231 | H33 | 86.2 | MET235 | H13 | 87.0 |
| 7 | MET235 | H13 | 85.8 | LEU231 | H39 | 87.0 |
| 8 | LEU231 | H41 | 84.7 | LEU231 | H41 | 86.4 |
| 9 | LEU231 | H40 | 84.4 | LEU231 | H36 | 85.2 |
| 10 | LEU231 | H39 | 84.2 | LEU231 | H37 | 84.7 |
| 11 | LEU231 | H38 | 81.9 | MET235 | H5 | 84.3 |
| 12 | LEU231 | H37 | 81.4 | LEU231 | H38 | 84.3 |
| 13 | MET235 | H5 | 81.2 | MET235 | H6 | 83.2 |
| 14 | LEU231 | H36 | 80.2 | LEU231 | H33 | 82.1 |
| 15 | THR239 | H16 | 79.9 | MET235 | H14 | 80.3 |
| 16 | MET235 | H18 | 77.8 | MET235 | H31 | 78.8 |
| 17 | MET235 | H14 | 77.7 | MET235 | H30 | 77.8 |
| 18 | MET235 | H19 | 76.3 | MET235 | H32 | 77.6 |
| 19 | MET235 | H31 | 75.1 | MET235 | H8 | 76.9 |
| 20 | MET235 | H11 | 75.1 | LEU231 | H49 | 75.5 |

**Table 3:** Top 10 BVM/CTD-SP1 molecular interactions involving CTD-SP1 lysine residues, with and without IP_6_. The contact occupancy measures how frequently the corresponding contact was formed in the simulation. A contact is defined when the distance between two groups of atoms is smaller than 3.0 Å.

| BVM-CTDSP1 interactions |  |  |  | BVM-IP<sub>6</sub>-CTDSP1 interactions |  |  |
| --- | --- | --- | --- | --- | --- | --- |
|  | CA residues | BVM atoms | Occupancies (%) | CA residues | BVM atoms | Occupancies (%) |
| 1 | LYS227 | C36 | 69.9 | LYS227 | C36 | 54.0 |
| 2 | LYS227 | O6 | 57.7 | LYS227 | O6 | 46.5 |
| 3 | LYS227 | O5 | 55.5 | LYS227 | O5 | 45.3 |
| 4 | LYS227 | H55 | 23.6 | LYS227 | H54 | 28.2 |
| 5 | LYS227 | H51 | 22.9 | LYS227 | H53 | 24.2 |
| 6 | LYS227 | H52 | 21.3 | LYS227 | H55 | 19.8 |
| 7 | LYS227 | H53 | 20.7 | LYS227 | H50 | 16.3 |
| 8 | LYS227 | H54 | 20.3 | LYS227 | H52 | 15.6 |
| 9 | LYS227 | H50 | 19.1 | LYS227 | H51 | 15.0 |
| 10 | LYS227 | H48 | 9.0 | LYS227 | H49 | 12.4 |

**Supplementary Figure 1:**
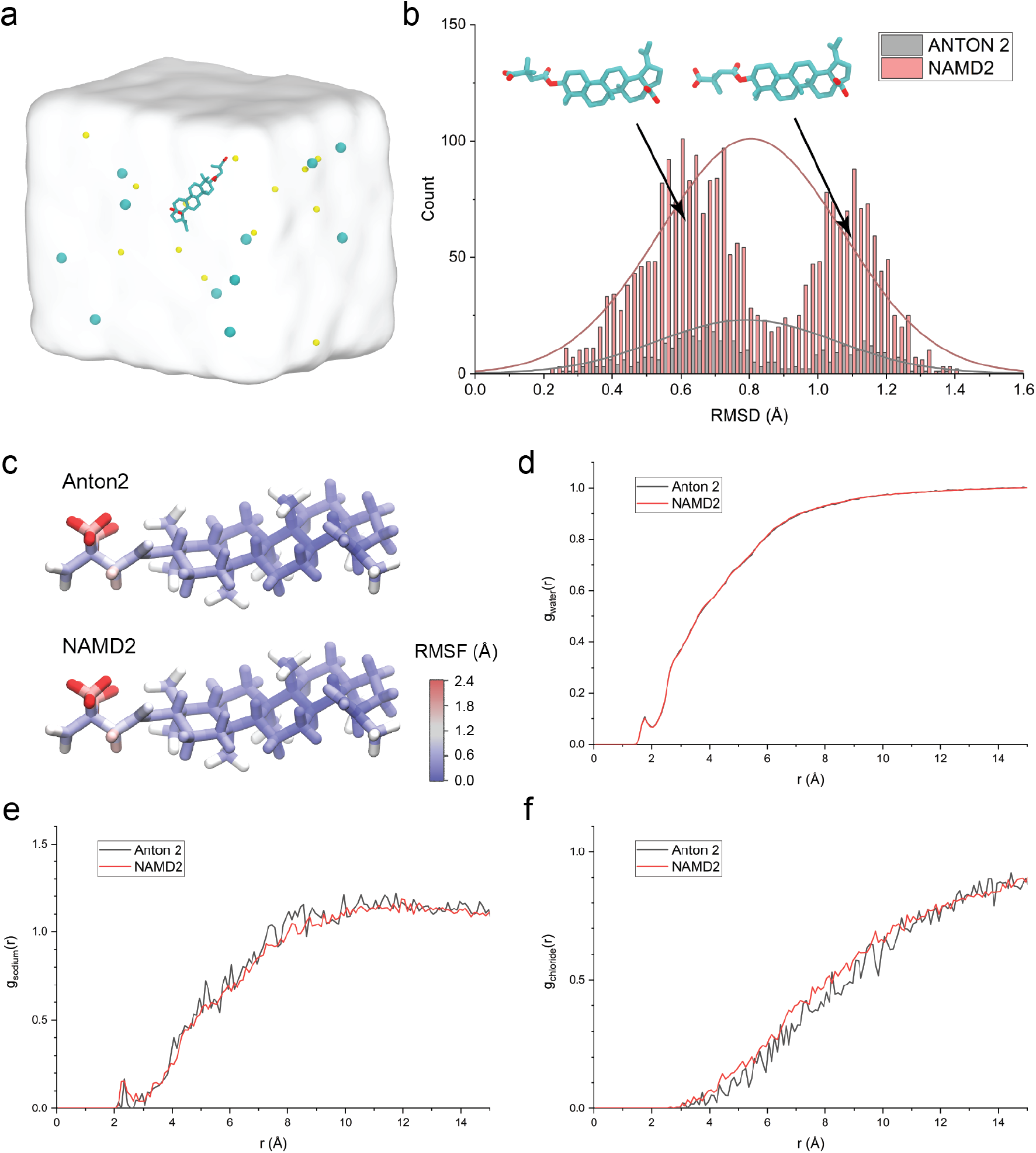
Dynamics of the solvated BVM molecule from NAMD2 and Anton 2 simulations. (a) A model of BVM in water box with 150 mM NaCl. (b) The distribution of the BVM RMSD in MD simulations. The initial conformation of BVM was used as the reference to calculate the RMSD. Two representative structures of two BVM RMSD clusters identified from MD simulations are illustrated as insets. (c) RMSF of individual atoms in BVM molecules from Anton2 and NAMD simulations. (d), (e) and (f) are the BVM-water, BVM-Na^+^ ion and BVM-Cl^-^ ion radial distribution functions calculated from MD simulations, respectively.

**Supplementary Figure 2:**
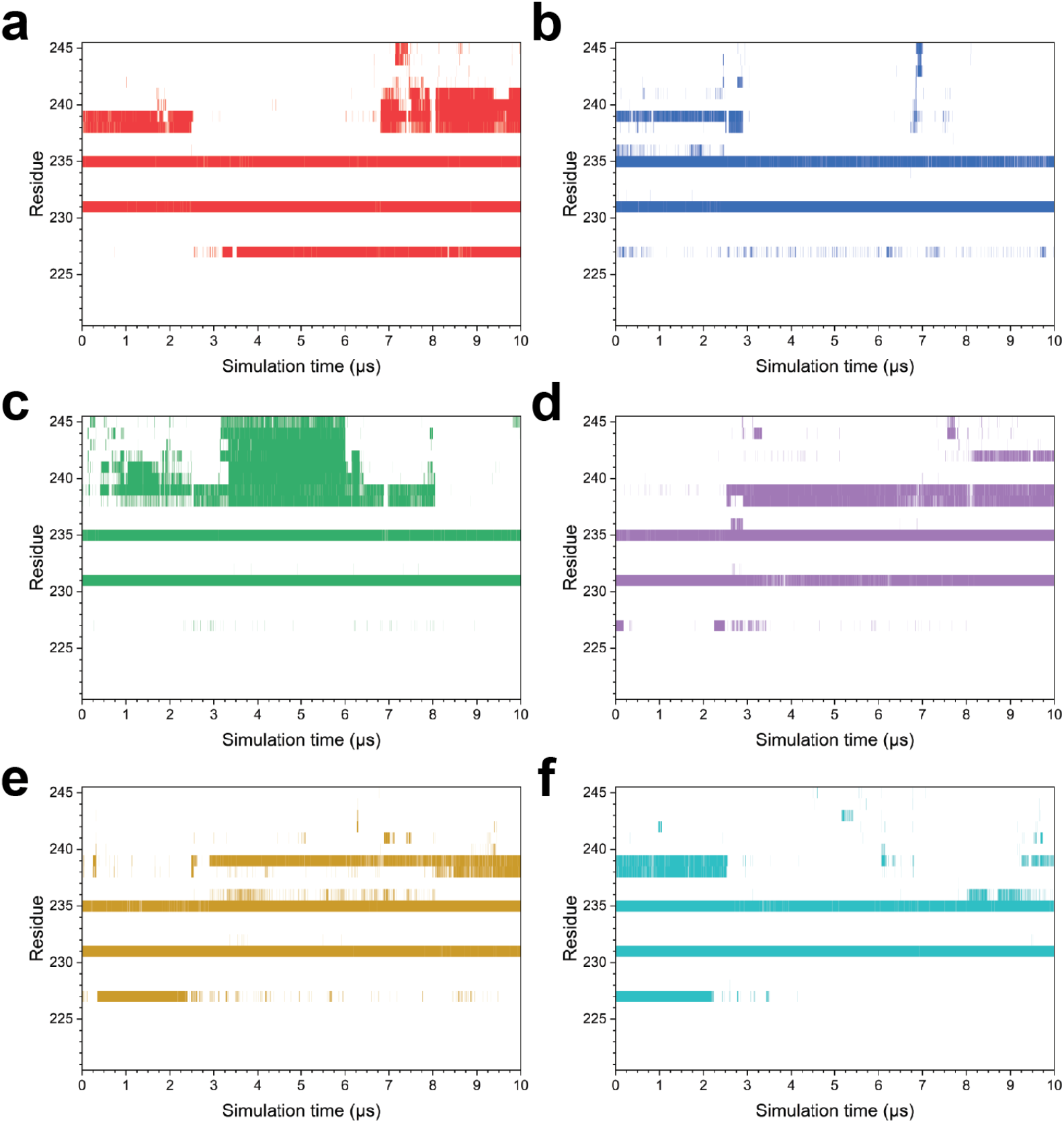
The molecular interactions between BVM and SP1 domain in the CTDSP1 hexamer. (a), (b), (c), (d), (e) and (f) BVM-Helix interactions from chain 1, 2, 3, 4, 5 and 6 in CTDSP1 hexamer over 10 *µ*s MD simulations.

**Supplementary Figure 3:**
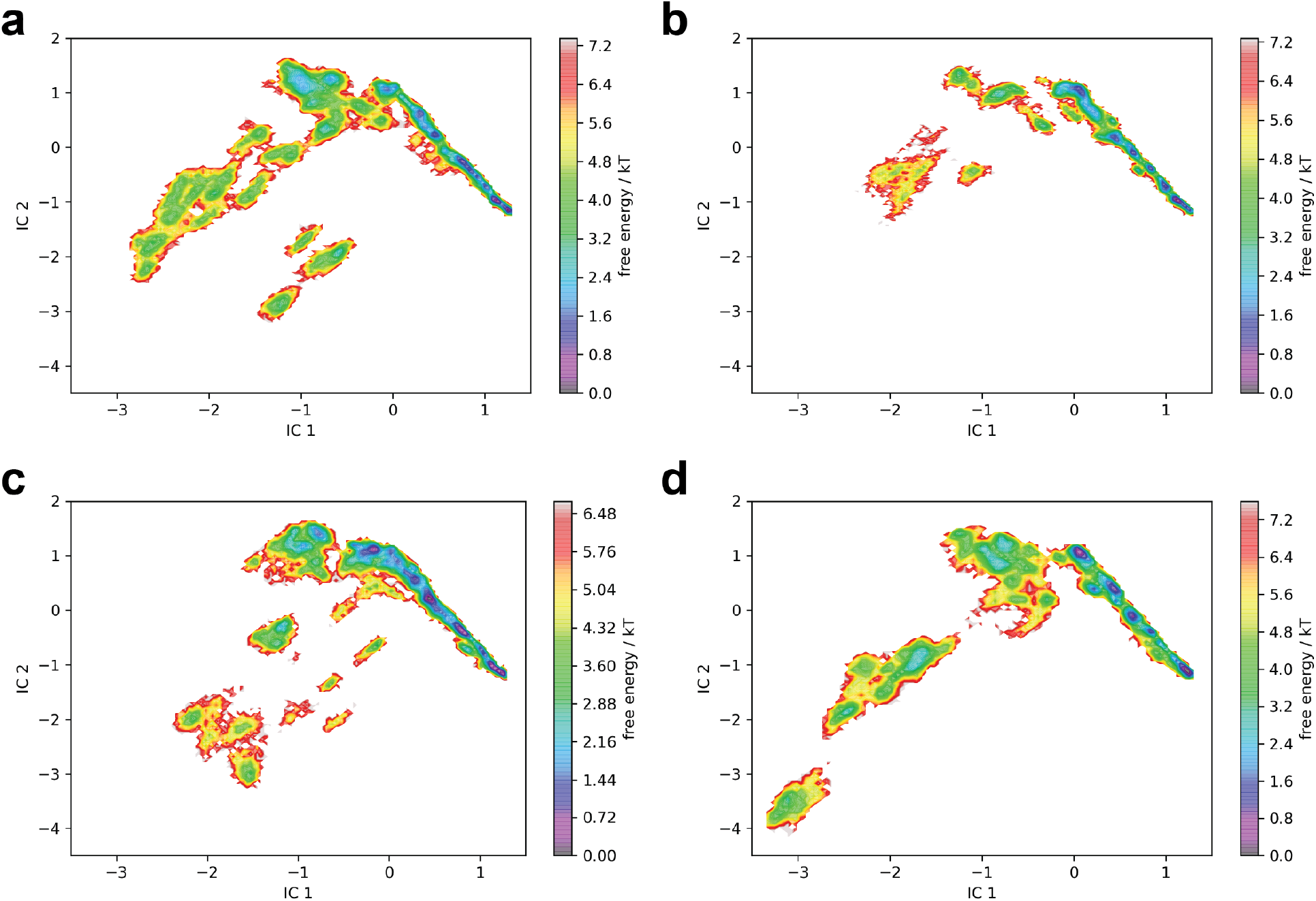
Transition path ensembles (TPEs) of SP1 domain present in CTD-SP1 hexamers for (a) apo, (b) IP6 bound, (c) BVM bound, and (d) IP6/BVM bound states. Comparison between the TPEs for bound and unbound BVM shows that binding of BVM shifts the TPE for the helix-coil transition on the CTD region while maintaining the TPE in the SP1 region. Binding of IP_6_ alters the TPE of the CTD region while preserving the dynamics of the SP1 region. The helix-coil TPE of the CTD region of the SP1 domain is modulated by IP_6_ and BVM binding while the dynamics of the SP1 subdomain is unaltered by any of these two molecules.

**Supplementary Figure 4:**
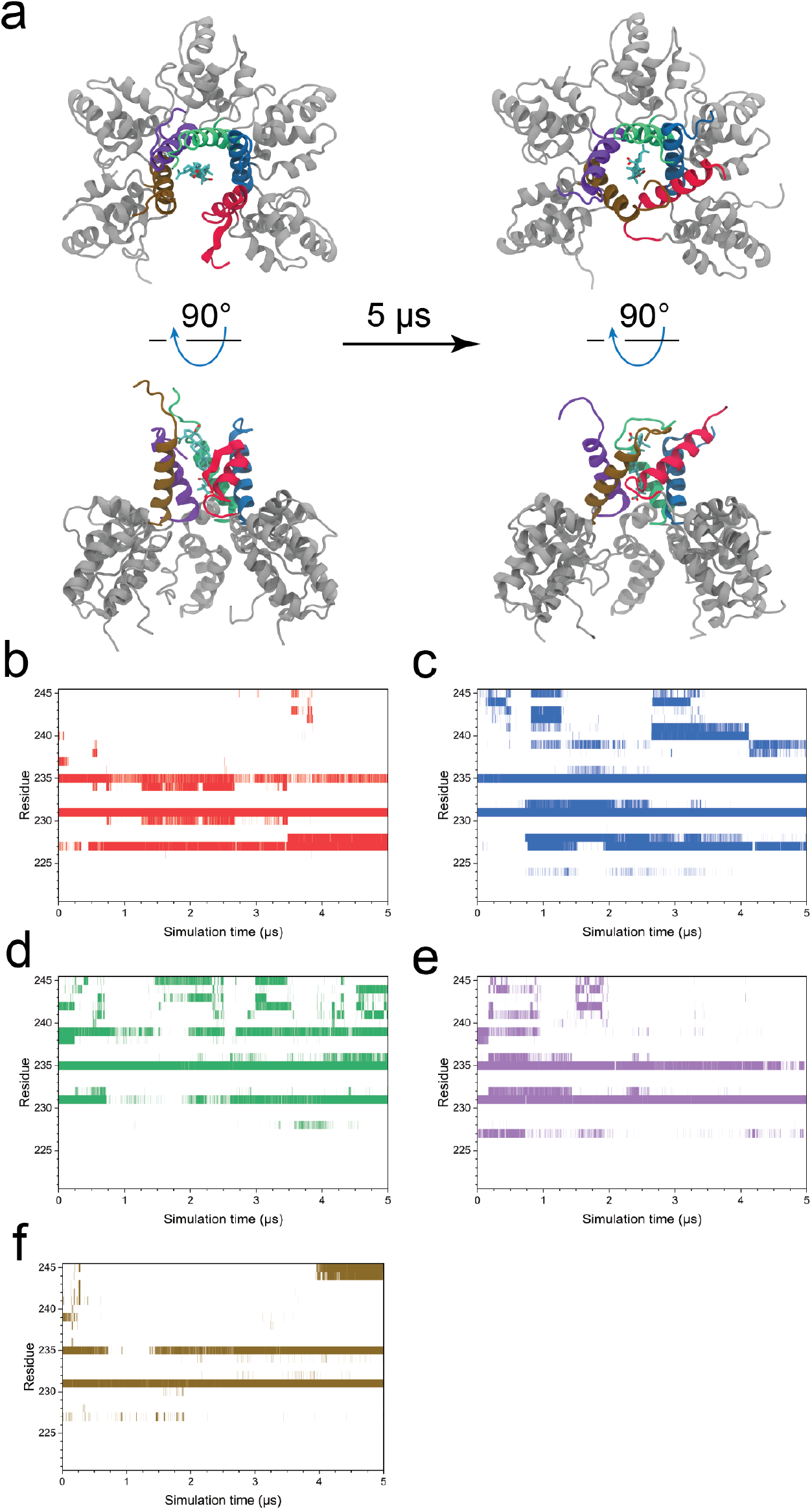
BVM/CTD-SP1 pentameric interactions. (a) The structures of pentameric CTD-SP1 and BVM complex before and after MD simulation. (b)-(f) Heat maps show the residues in the SP1 domain interacting with the BVM molecule during the simulation. SP1 domain shown in (a) are distinguished by different colors, which correspond to the colors in the heat maps.

**Supplementary Figure 5:**
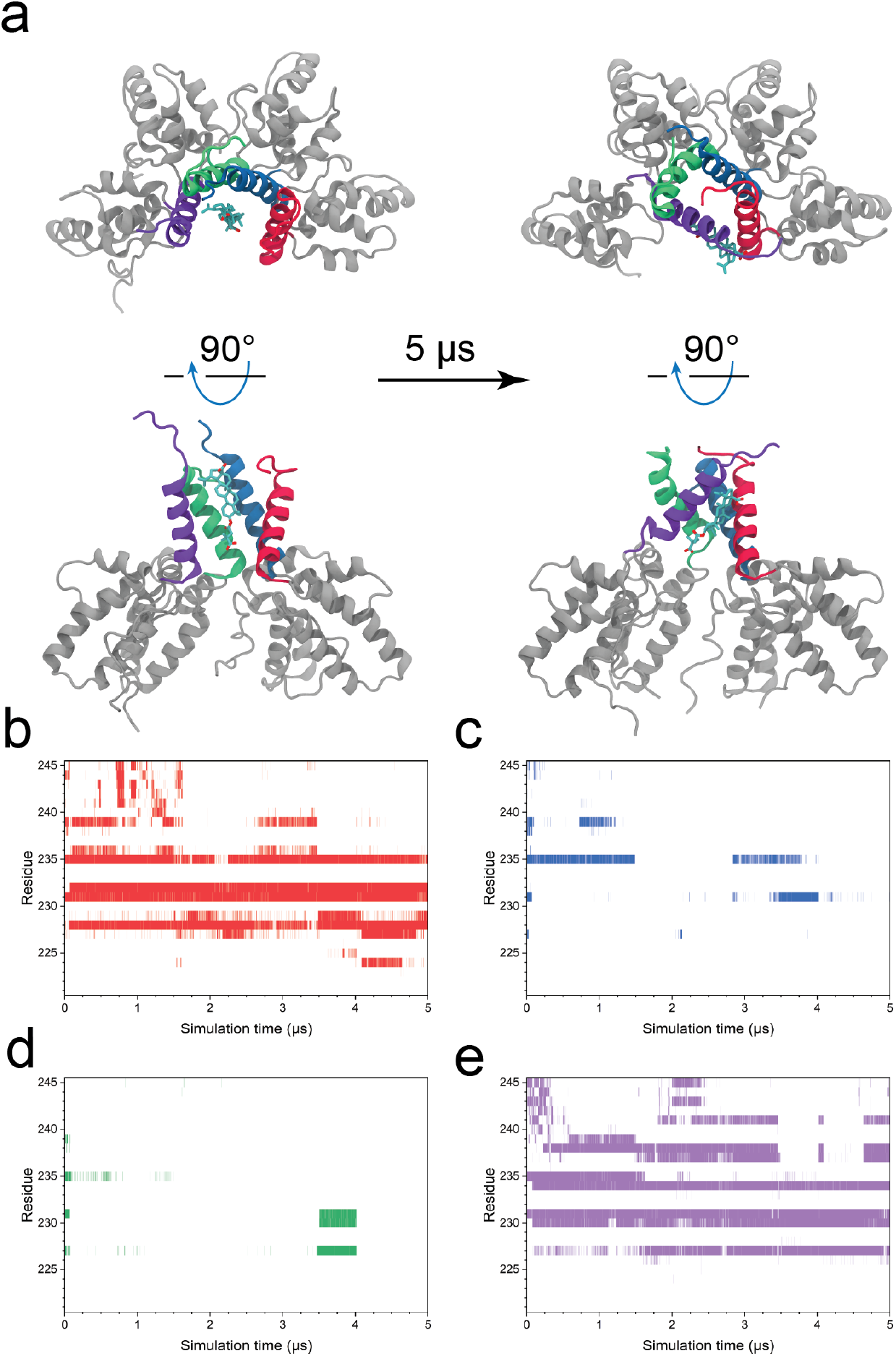
BVM/CTD-SP1 tetramer interactions. (a) The structures of tetrameric CTD-SP1 and BVM complex before and after MD simulation. (b)-(e) Heat maps show the residues in the SP1 domain interacting with the BVM molecule during the simulation. SP1 domain shown in (a) are distinguished by different colors, which correspond to the colors in the heat maps.

**Supplementary Figure 6:**
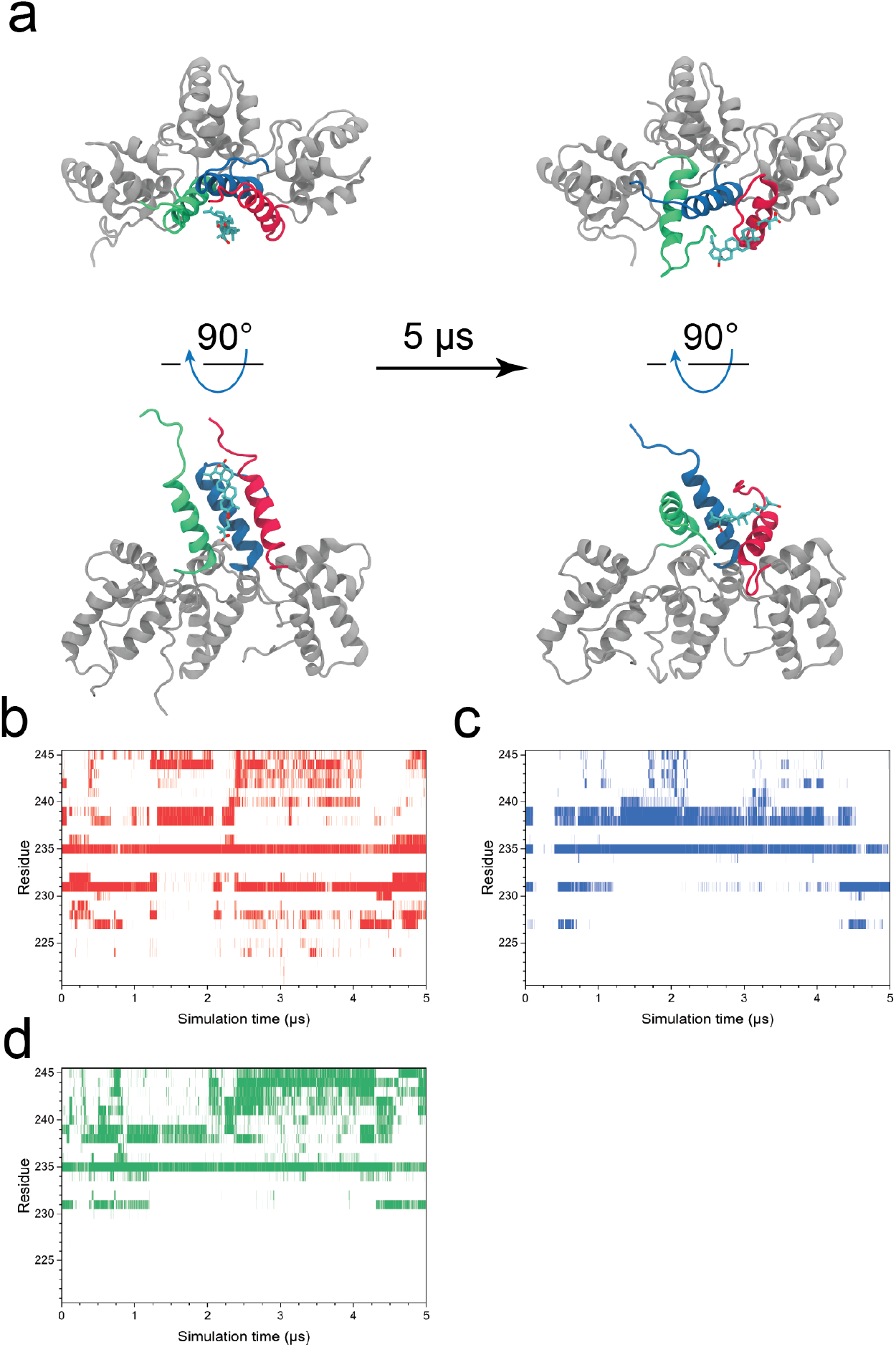
BVM/CTD-SP1 trimeric interactions. BVM-CTDSP1 3mer interactions. (a) The structures of trimeric CTD-SP1 and BVM complex before and after MD simulation. (b)-(d) Heat maps show the residues in the SP1 domain interacting with the BVM molecule during the simulation. SP1 domain shown in (a) is distinguished by different colors, which correspond to the colors in the heat maps.

**Supplementary Figure 7:**
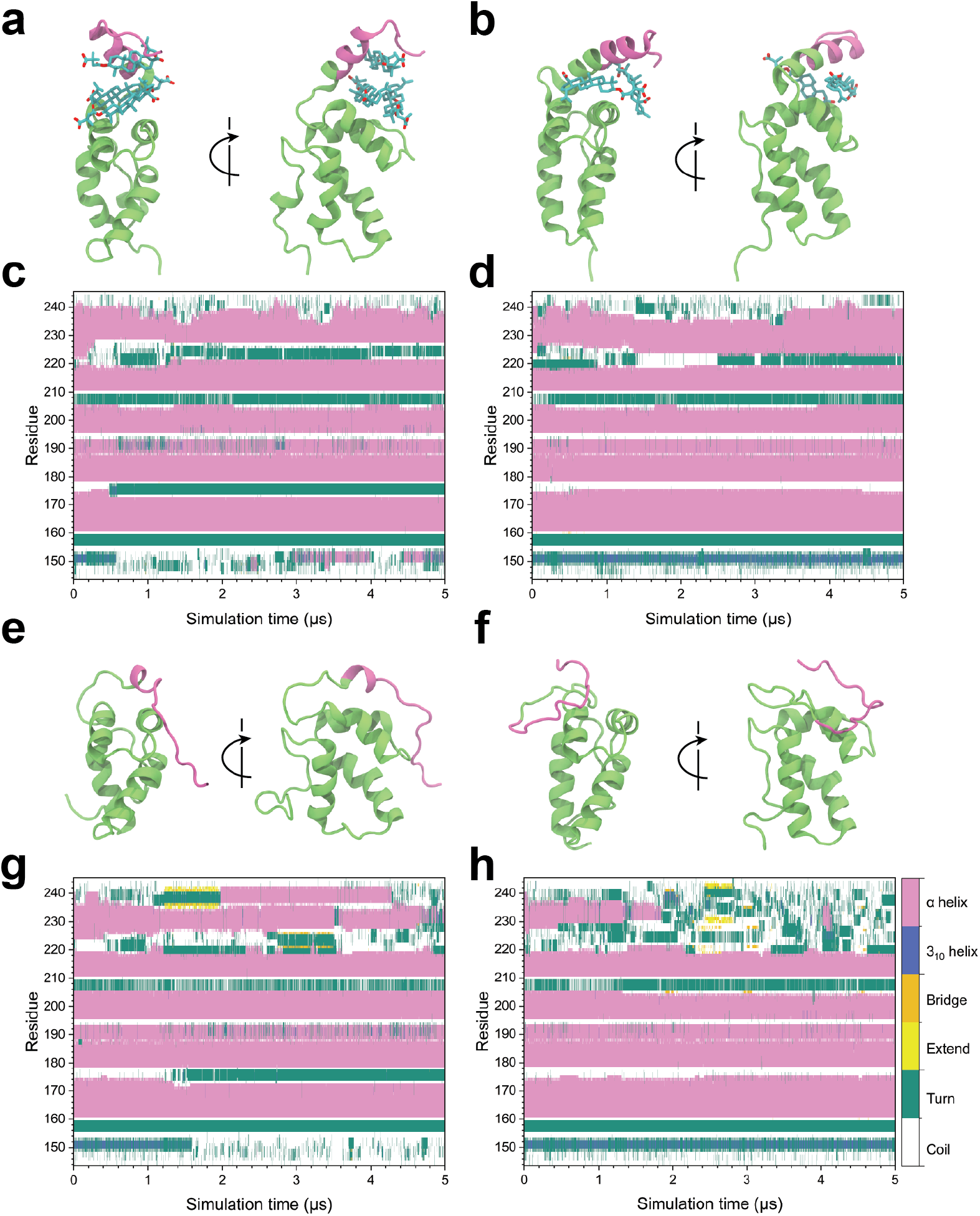
BVM-CTDSP1 monomer interactions stabilize the SP1 domain. (a) and (b) BVM-CTDSP1 monomer system from two independent 5 *µ*s flooding MD simulation run #1 and run #2, respectively. The BVM-CTDSP1 monomer system includes five free BVM molecules and one CTDSP1 monomer in explicit solvent. After simulations, only those BVM interacting with the SP1 domain are shown in the structures. The secondary structure transitions of the CTDSP1 from two simulations are shown in (c) and (d). (e) and (f) CTDSP1 monomer system from two independent 5 *µ*s MD simulation run #1 and run #2, respectively. The CTDSP1 monomer system, has a similar size as the BVM-CTDSP1 monomer system, contains one CTDSP1 monomer and no BVM molecules. The secondary structure transitions from these two simulations are displayed in (g) and (h), respectively. The color bar on the right side of panel (h) indicates the correspondent secondary structures in the heat maps.

**Supplementary Figure 8:**
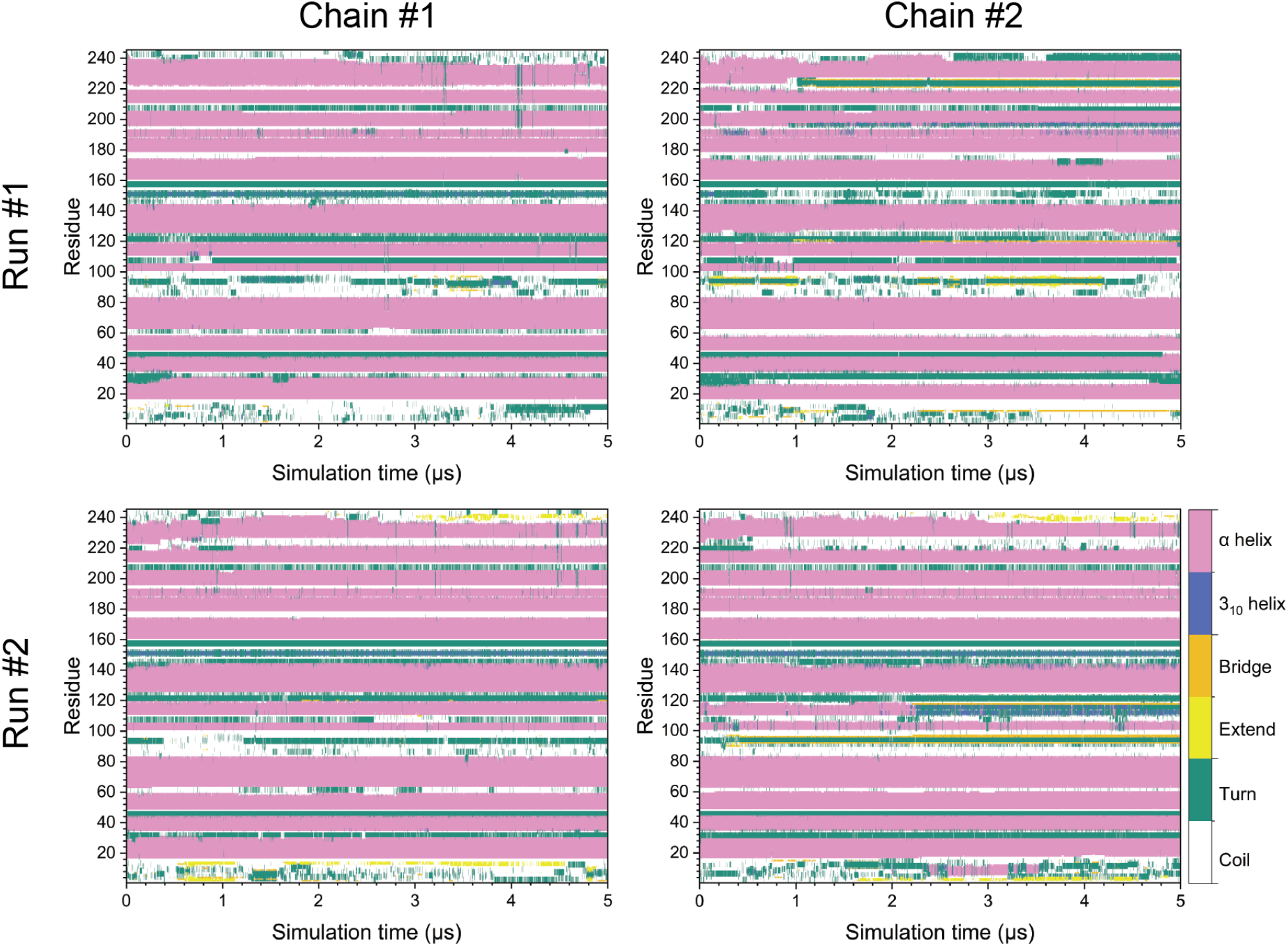
The secondary structures of CASP1 dimer systems from two independent BVM flooding MD simulations. Figures in the first and second row show results from two different simulations. The color bar in the bottom right indicates the correspondent secondary structures in the heat maps.

